# Capture and Release of Minke Whales Offers New Research Opportunities Including Measurements of Mysticete Hearing

**DOI:** 10.1101/2024.02.02.577929

**Authors:** Lars Kleivane, Petter H Kvadsheim, Anna Victoria Pyne Vinje, Jason Mulsow, Rolf Arne Ølberg, Jonas Teilmann, Craig Harms, Dorian Houser

## Abstract

Knowledge about species-specific hearing is vital to assessing how anthropogenic noise impacts marine mammals. Unfortunately, no empirical audiogram exists for any mysticete whale. We therefore developed a catch-and-release method to assess hearing in a small mysticete, the minke whale (*Balaenoptera acutorostrata*). Stationary lead nets were placed to intercept migratory routes and direct animals into an ocean basin enclosed by nets and islets, while another net was pulled across the entrance once a whale entered the basin. The whales were then slowly corralled into a modified aquaculture pen using a net suspended between two boats. Subsequently, the water volume available to the whales was gradually reduced by raising the pen net by hand until the whales were secured in a “hammock” between the floating pen ring and a raft. From the raft, researchers could access the whales to monitor their health, apply instruments for hearing tests, or other research objectives, and attach tags to monitor the movements and diving behavior of the whale post-release. The method is a slow and controlled procedure, allowing continuous monitoring, and quick release of the whales, if needed. In the first three field seasons employing the method, three animals were caught for research procedures. Initial hearing measurements using auditory evoked potentials were successfully completed. After release the animals resumed migration and dive behavior were considered normal. Our observations demonstrate that minke whales can be safely guided via moored net barriers, corralled into an aquaculture pen, and safely handled for research purposes, before being released back into the wild.

## Introduction

Cetaceans are divided into the toothed whales (odontocetes) and baleen whales (mysticetes) and are distinguished by fundamental differences in functional morphology, sensory physiology, dive adaptations and feeding behavior (Reynolds III & Rommel, 1999). Several odontocete species are kept under human care in aquariums or dedicated research facilities, where, under strong ethical scrutiny, researchers are allowed to collect blood, take tissue biopsies, or attach instruments to study their physiology. Research with odontocetes under human care has increased biological knowledge about these animals and has subsequently led to better management of wild populations. Primarily due to their size, mysticetes are not kept under human care and are only briefly held during stranding or entanglement responses. Some behavioral and physiological data may be acquired by the deployment of tags on mysticetes. Such technology is limited, and much of our current understanding of mysticete physiology is derived from postmortem examinations (Goldbogen et al., 2015), or extrapolation from odontocetes or pinnipeds. As a result, the validity of our understanding remains in question (e.g., with respect to diving adaptations (Fahlman, 2012; Hooker et al., 2012), hearing (Cranford & Krysl, 2015; Southall et al., 2019), or thermoregulation and energetics (Lavigne et al,. 1986)).

Hearing in mysticetes is predicted from anatomical models (Houser et al., 2001; Parks et al., 2007; Cranford & Krysl, 2015; Tubelli et al., 2018), the frequency range of mysticete vocalizations (e.g., Schevill & Watkins, 1972; Clark & Johnsen, 1984; Cummings & Thompson, 1994; Thompson et al., 1986), and behavioral responses of mysticetes to incidental and intentional noise exposures (e.g., Todd et al., 1992; Goldbogen et al., 2013; Curé et al., 2015; Sivle et al., 2016; Kvadsheim et al., 2017, Boisseau et al., 2021). Since no empirical measure of hearing range or sensitivity has been made in a mysticete whale outside of an unsuccessful attempt by Ridgway & Carder (2001) to measure the hearing of a stranded gray whale (*Eschrichtius robustus*) calf, direct measurements of mysticete hearing are needed to validate models and inform knowledge gaps identified by scientists (e.g., Southall et al., 2019), marine noise polluters (IOGP, 2018) and noise regulators (NOAA, 2018). Since mysticete whales are not kept under human care and behavioral hearing tests (the most accurate measure of audition) are not possible to perform, the most likely method of directly measuring hearing in a mysticete is through auditory evoked potential (AEP) methods. These methods have been broadly used and validated on odontocete species (e.g., Szymanski et al., 1999; Yuen et al., 2005; Finneran & Houser, 2006; Houser & Finneran, 2006a; Houser & Finneran, 2006b; Finneran et al., 2009; Ruser et al., 2016; Houser et al., 2022), and have great promise for use in smaller mysticete species. Nevertheless, the method requires that subject animals are temporarily caught and controlled for the hearing tests.

Previous attempts to catch live mysticete whales for scientific purposes have had limited success (Vinje, 2022). In the late 1950s, there was an unsuccessful attempt to capture a minke whale (*Balaenoptera acutorostrata)* calf for an aquarium. The calf was lassoed by the tail and hoisted onboard a research vessel, but died upon arrival at the aquarium (Norris & Prescott, 1961). Another aquarium succeeded in obtaining three minke whales between 1930 and 1954 (Kimura & Nemoto, 1956). A fishing net between two boats was used to capture the whales, which were then towed ashore. The animals escaped or died after a few weeks, and scientific measurements were limited to recordings of swim behavior and respiration rate (Kimura & Nemoto, 1956). In the 1960’s and 1970’s, SeaWorld captured two gray whales (*Eschrichthius robustus*) using a superficial harpoon followed by netting or a tail noose deployed from a fishing vessel (Norris & Gentry, 1974). The first whale died within two months from lung injury and pneumonia caused by the capture technique (Wahrenbrock et al., 1974). The second whale was kept in captivity for a year before it was released (Norris & Gentry, 1974). Respiratory physiology, circulatory physiology, energetics, vocalization, hematology, and feeding behavior were investigated in these two whales (Curran & Asher, 1974; Duffield 1974; Evans, 1974; Fish et al., 1974; Gilmartin et al., 1974; Leatherwood, 1974; Mattson, 1974; Medway, 1974; Norris & Gentry, 1974; Ray & Schevill, 1974; Smith & Wahrenbrock, 1974; Wahrenbrock et al., 1974, Zettner, 1974), but no direct measurements of hearing were made. Opportunistic scientific use of mysticete whales accidentally trapped in fishing gear (Winn et al., 1979), in natural enclosures (Beamish, 1979), or live stranded (Edds et al., 1993; Priddel & Wheeler 1998; Reidarson et al., 2001; Sumich, 2001) have occurred, but again no direct measurements of hearing have been made. The most recent attempt to intentionally live capture mysticete whales for research was specifically conducted to collect AEP audiograms. The effort took place in Iceland in 2007 (IOGP JIP, 2022). An international research team used a modified herring purse seine net to entrap minke whales and had a pontoon boat fitted with a stretcher to restrain a caught whale (J. Teilmann, personal observations). Eighty-one minke whales were observed during 11 days at sea, and a capture net was set 4 times with no successful catch. The nets were too heavy, took too long to set around the whales, and the animals consequently escaped the capture attempts.

Development of a safe and ethically acceptable method to live-catch and temporarily constrain mysticete whales, without compromising animal health and welfare, will not only provide opportunities to measure hearing, but also to study other aspects of mysticete whale biology, including sensory and respiratory physiology, energetics and instrumentation for behavioral and physiological studies of free-ranging whales. The objective of the effort reported here was to develop a safe catch-and-release method for minke whales for the principal purpose of obtaining an AEP audiogram. Minke AEP hearing tests will provide the first direct measurement of hearing in a mysticete whale, but the catch-and-release method could have broader potential for studying mysticetes in the wild.

## Method

The methods described here are meant to target adolescent minke whales (*Balaenoptera acutorostrata*), partly because of the practical challenges of handling larger animals, but also because AEPs are more difficult to measure in larger animals due to their relatively smaller brain-to-body mass ratios. Minke whales arrive along the Norwegian coast for their seasonal feeding migration in April-June, and a relatively high number of younger animals make a detour into Vestfjorden on their way northwards to the rich feeding grounds in the Barents Sea (Christensen & Rørvik, 1981; Heide-Jørgensen et al., 2001). The field site was located off Stamsund in Lofoten (Norway). The site was chosen after numerous interviews with local fishermen about whale presence in this region, as well as prior experience during whale research in the same area (Walløe et al., 1995), during which minke whales were observed travelling westwards along the Lofoten peninsula, close to and between the numerous Lofoten islands (Figure 1). June, which has generally favorable weather conditions and high whales density, was chosen as the optimal catch period. Catch attempts have been made in three field seasons so far; June 2021, 2022 and 2023.

**Figure 1.**
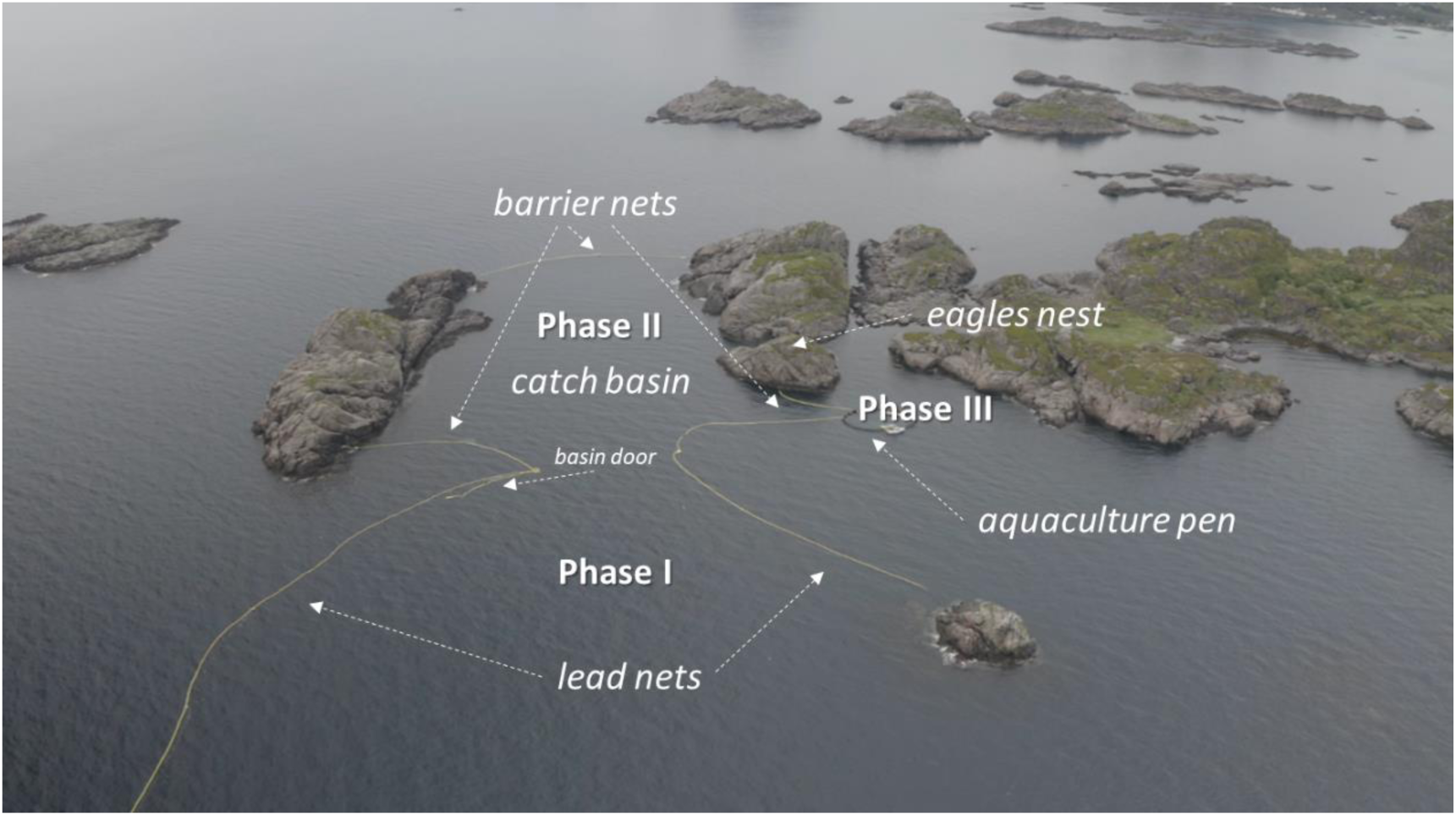
Drone picture of the catch and release site. The catch process is divided into three phases. Phase I is the catch phase, which ends when the netted basin door behind the whale is closed and the whale is contained in the catch basin. Phase II is the corralling phase, which ends when the door of the aquaculture pen is closed behind the whale. Phase III is the final phase where the whale is placed and held in a net hammock and ends when the whale is released back to the wild. Whales are monitored from the “Eagle’s Nest” and/or from boats docked at the aquaculture pen from the first sighting in Phase I until released at the end of Phase III (Figure 5). Photo: E. Wang-Naveen, FFI.

### The Catch and Release Site (CARS)

The catch-and-release site (CARS) consisted of 1790 m of medium-meshed (65-78 mm mesh size) purse seine nets ranging from 20-55 m depth. Net depths were adapted to the bathymetry profile of the site. Anchored lead nets were set to intercept the migration route and direct the animals between the two islets of Kvannholmen and Æsøya (Figure 1). On both the west and east side of these islets, barrier nets blocked the gap between them, to form a large basin (280 m long, 160 m wide and 20-30m deep) with a volume of about 1x10^6^ m^3^. The barrier net blocking the west side of the basin (*A-net*) was anchored to the islets. The barrier nets blocking the east side of the basin (*B-nets*) contained a 40-m wide opening (CARS-door) and were anchored to the islets and to the aquaculture pen (Figure 2). The circular aquaculture pen was 90-m in circumference and contained both inner and outer nets affixed around its entire circumference. Anchored on the south side of the CARS-door, lead nets (*D-nets*, 1100 m in length, 45-55 m in depth) stretched eastwards to a shallow grounding point (Brusen) where the net was anchored. On the north side of the basin opening, a 160-m lead net (*C-net*, 25 m deep) was anchored and stretched eastward and attached to another small islet, Ausa (Figure 2). A single unanchored net (*E-net*, 100 m in length, 20 m in depth) was tethered along the A-net until it was used for corralling a whale towards the aquaculture pen. Between field seasons slight modifications were made to the net configuration. The main adjustment was between year 2 and 3 where the outer part of the D-net (D2) was removed, and the inner part (D1) was bent to the northeast and attached to the islet Flatskjaeret (Figure 2). This modification was based on experiences from the first two years to increase the catch rate.

**Figure 2.**
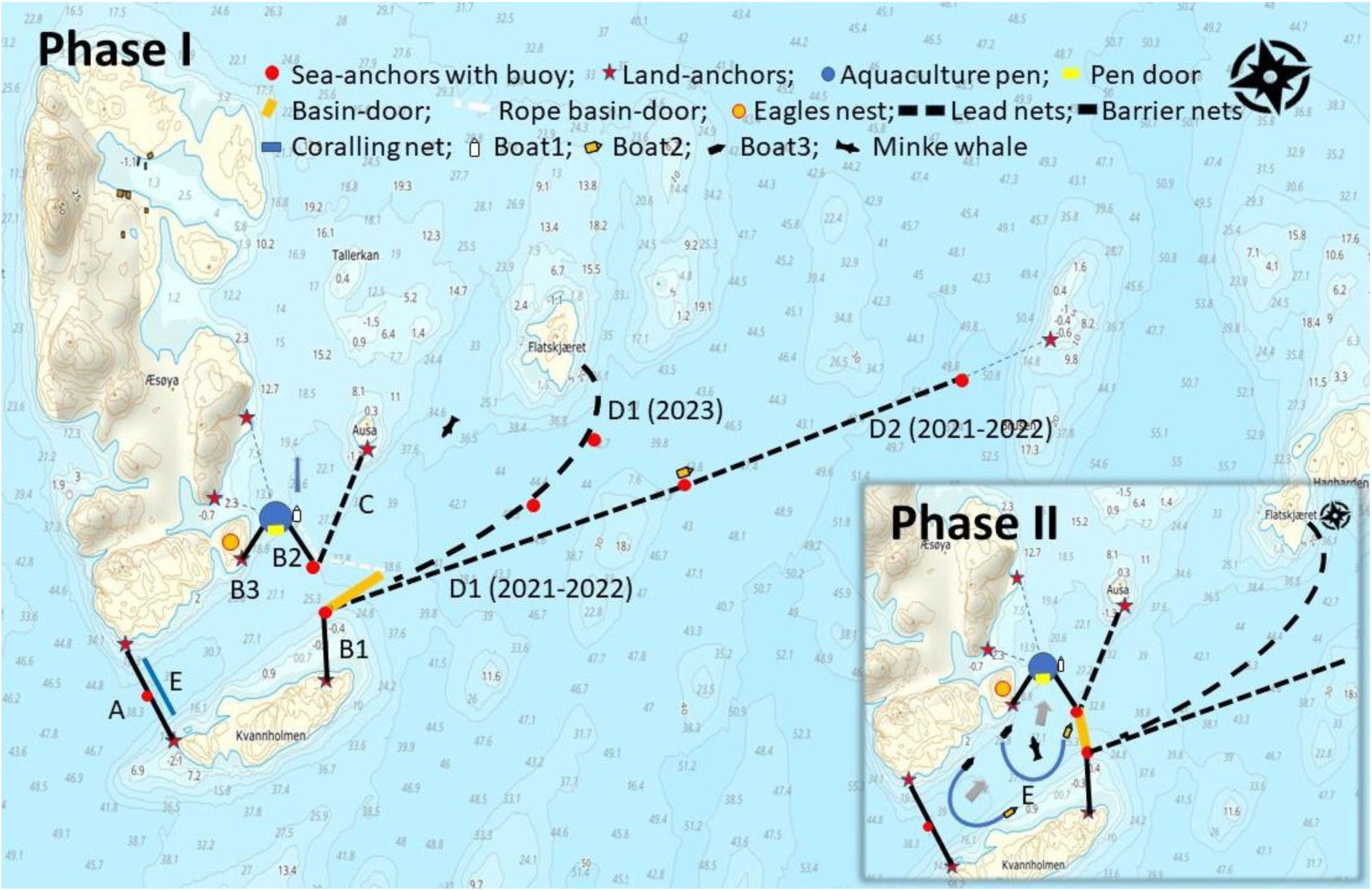
The trap design for live-catch and release of minke whales in Lofoten (Norway). The map shows details of the placement of nets (A:160 m, B1:120 m including 40 m trapdoor, B2:100 m, B3:50 m, C:160 m, D1:600 m, D2:500 m; E:100 m), anchors, 90 m aquaculture pen, observer platforms at the Eagle’s Nest and boats (Boat 1 closing the door and boat 2 patrolling the area). The configuration of the lead nets (C and D) changed somewhat between years (Figure 4). Insert: Map showing the corralling path of a minke whale from the catch basin into the aquaculture pen in phase II of the catch process. The 100-m long E net is pulled between two boats from the A net eastwards towards the door of the aquaculture pen.

To provide resilience to tidal and coastal currents in the CARS, all nets were heavily weighted with two bottom lead lines, the main bottom line had 8.4 kg/m weight and the secondary weight line above it had 2.0 kg/m weight. The top of the net had 28 kg/m of buoyancy (four 7-liter buoys/meter). The tidal variation is up to 3 meters in the area; thus, all barrier nets (A, B1, B2, B3 in figure 2) were constructed with a tidal skirt with two weight lines vertically separated by 3 m in order to reach the bottom at high tide, while allowing them to be held tight without excessive slack at low tide. All nets were constructed at Mørenot AS (Ålesund, Norway) from herring and mackerel fishery purse seine nets. They were constructed with 210D nylon netting material of 144-178 PLY. The nets were cut into bar meshes before mounting to obtain a stable depth and length and to reduce drag on the nets by allowing water flow through them. The lead and barrier nets were deployed off a large purse seine fishing vessel (>20m long) with a triplex net handling system and >50 m^3^ net loading capacity. Several trips were required to deploy all of the nets used in the CARS construction (Figure 1). Between years, all nets and lines were stored in either flexible bulk bags (big-bags with 6-8 m^3^ capacity) or in 12.2 m steel containers.

The aquaculture pen consisted of double, heavy-duty polyethylene floating pen rings (315 mm diameter) with a stanchion thick-walled (125 mm diameter) handrail. The pen had a buoyancy of more than 10 tons. Two aquaculture nets were used during the catch; one outer net with a door constructed in it (10 x 15 m), and an intact inner net without openings. The aquaculture pen nets had a mesh size of 31 mm and were composed of 1.7 mm diameter nylon twine. Top, waterline, bottom, vertical and horizontal ropes of 18 mm braided Danline were sewn into the nets for structural support and hooks were mounted into the handrail for attaching excess during the constraining phase. Each net was made as a cylinder module with a straight wall to 15 m of depth, and was then extended further 7 m in a coned shape that ended in a weighted center (30 kg). The net had a volume of more than 10,000 m^3^. Due to strong tidal currents, the net interiors were weighted with ten rounded, 30-kg weights suspended from adjustable ropes that were attached to the pen ring. Both aquaculture nets also had 22 vertical ropes tied to the bottom of the net and attached to the top of the ring at even spacing. The ropes were used along with the hooks in the handrail to adjust the depth of the inner net and to tie up the ends of the pen door when open. A raft (8 x 3 m) was constructed for this study and placed inside the aquaculture pen and used during handling of the whale in the final phase of the experiment (Figure 3).

**Figure 3.**
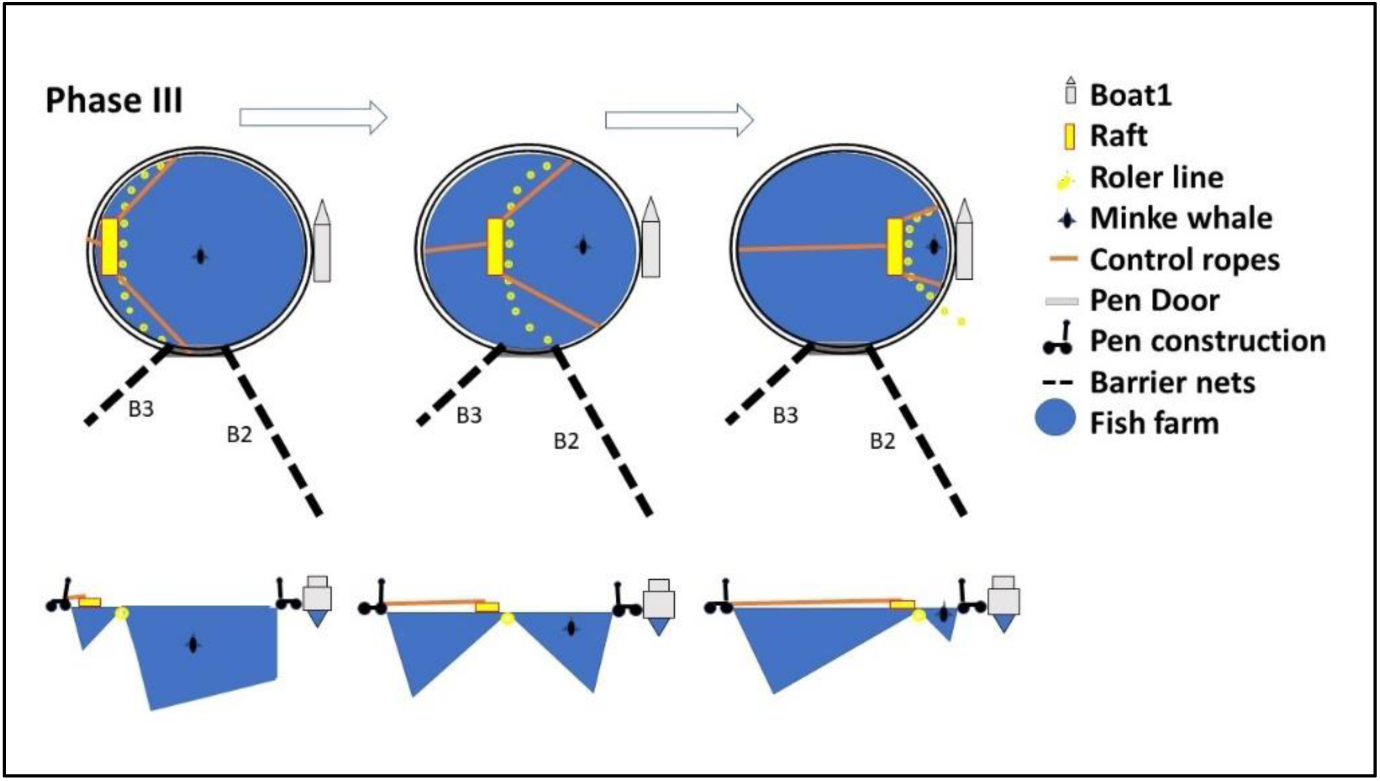
Phase III - getting the whale into the net hammock. The water volume around the whale is progressively reduced using a floating roller line (view from above in top row, profile in bottom row; progression occurs from left to right). Crew on the raft access the whale for health monitoring (e.g., ECG electrode attachment, blood draws, etc.) or to perform other investigations (e.g., AEP hearing measurements, tagging). Crew on both sides help to keep the whale stabilized in the hammock. If necessary, the whale can be quickly released back into the aquaculture pen by pulling out the floating roller line and dropping the net.

Weather was a challenging factor at the field site; thus, the aquaculture pen was anchored to Æsøya so that it was sheltered from strong southwesterly winds. The aquaculture pen was acquired from a local aquaculture farm (IsQueen AS) and deployed by their 15 m dredger boat. The boat was also hired to adjust barrier and lead nets, and for the deployment of anchor moorings. The floating aquaculture pen ring and nets were secured with strong ropes (30 mm Mixed Dyneema-Polyester) to eye-bolts on land at either Æsøya, Kvannholmen or Ausa (Figure 2), in addition to five, 1200 kg moored anchors (Figure 2).

During whale catches, a boat (Boat #1; 8.8 m Halco Offshore with a Volvo Penta 250 HP engine) was docked facing northward at the aquaculture pen. A 200 m long braided rope (18 mm) was attached to the stern of the boat with the opposite end attached to the CARS-door (Figure 2). The rope was dark colored and sank to the bottom of the ocean floor to avoid being perceived by the whales. A second boat (Boat #2; a 4.9 m fiberglass boat with an outboard four stroke 90 HP engine) stayed either docked at the aquaculture pen next to Boat #1, or patrolled the catch area, depending on the situation and the weather.

### The Catch Process

The catch process was divided into three phases (Figure 1): Phase I – catching the whale in the basin; Phase II - observing and corralling the whale into the aquaculture pen; and Phase III - constraining the whale for instrumentation and experimentation.

#### Phase I; Catching the Whale

During June, northern Norway experiences 24 hours of sunlight, and the CARS was monitored for 20 hours of the day using a crew of 12 people (two ten-hour shifts with six people per shift). During each shift, whale lookouts were stationed at an observation point on a hill located on Æsøya (*Eagles Nest*, 18 m height), as well as from two boats. Whales were visually tracked, until they left the area of the CARS. Respiration rates were recorded and swimming behavior were scored as either calm and normal swimming (CNS), fast vigorous swimming (FVS), or spy hopping (SPY). The CARS was divided into 8 zones (0-7), primarily to be able to quickly communicate sighting positions (Table 1, Figure 4). As soon as a whale was sighted in the catch basin (Zone 0), all personnel were alerted via hand-held radio to close the entrance to the catch basin. Boat #1 then immediately motored northward, pulling on the line to close the catch basin door. Once shut, the crew on boat #2 secured the net door in the closed position with a line and quick-release carabiners.

**Figure 4.**
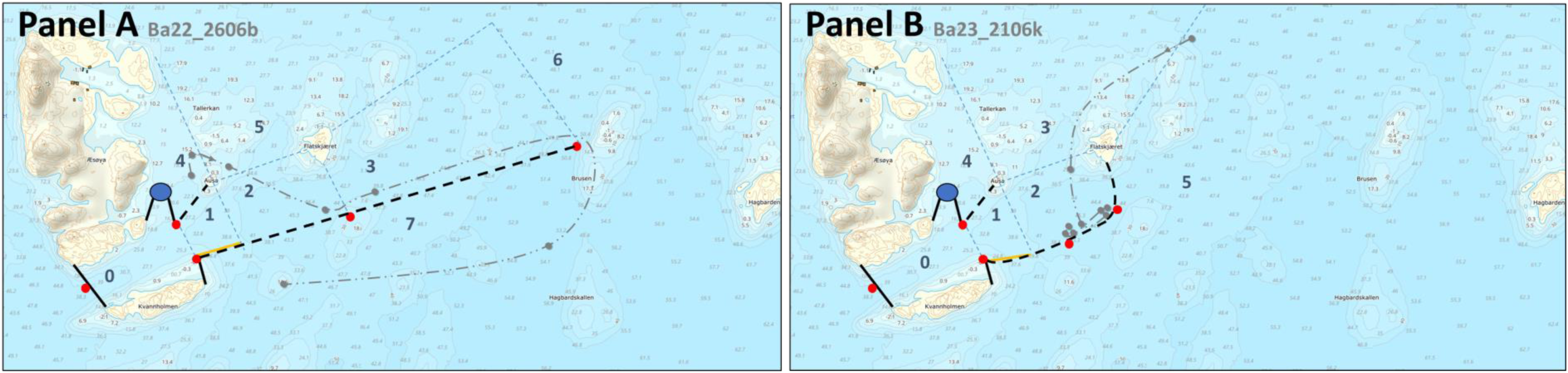
Example tracks of whales (light grey) approaching the CARS with the 2021/2022 net configuration (Panel A) and with the 2023 net configuration (Panel B). The CARS was divided into zones (0-7), primarily to enable rapid communication of whale sightings. The examples show the whales typically entering the CARS from the north (Table 1). With the 2021/2022 net configuration many whales escaped the CARS eastwards along the D nets (Panel A). When the net configuration was changed in 2023, catch rates increased from 1-2 animals/year in 2021/2022 to 7 animals caught in the catch basin (Zone 0) in 2023.

**Table 1.**
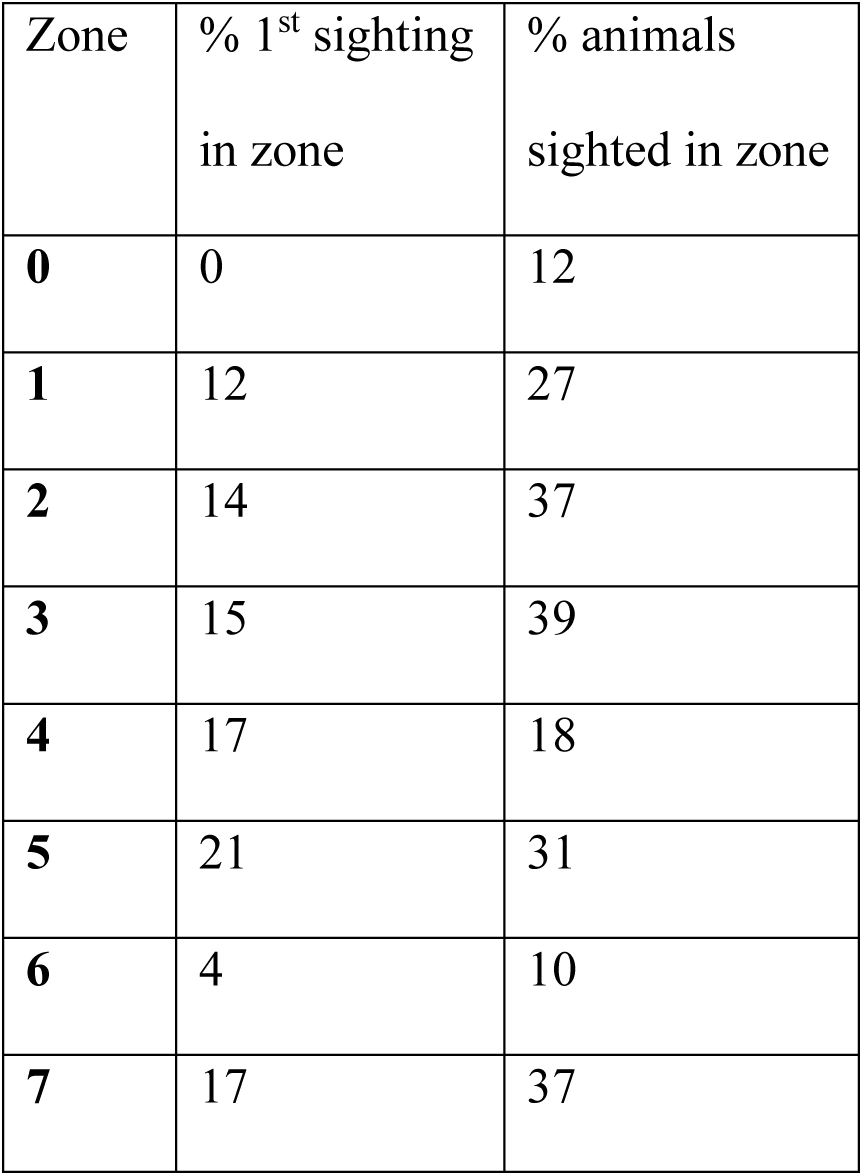
The percent of first sightings in the different zones of the CARS with the net configuration used in 2021 and 2022 (Figure 4A), and the percent of the minke whales tracked through the CARS sighted in each zone. Most animals approached from the north (38%, first sighted in zone 4 and 5), fewer approached from the east (19%, first sighted in zone 3 and 6), and 26% approached from an unknown direction (first sighted inside CARS, zone 1 and 2). Only 17% of the animals sighted passed outside of the CARS (first sighted in zone 7). Based on these data from 2021/2022 the configuration of the lead nets was modified in 2023 (see figure 2 and 4).

| Zone | % 1<sup>st</sup> sighting<br>in zone | % animals<br>sighted in zone |
| --- | --- | --- |
| **0** | 0 | 12 |
| **1** | 12 | 27 |
| **2** | 14 | 37 |
| **3** | 15 | 39 |
| **4** | 17 | 18 |
| **5** | 21 | 31 |
| **6** | 4 | 10 |
| **7** | 17 | 37 |

#### Phase II; Observation and Corralling of the Whale into the Aquaculture Pen

After successful containment of a minke whale in the catch basin, the swim behavior and respiration rate of the whale was monitored from Eagle’s Nest for a minimum of 2 hours in 10-min intervals (Figure 2). If the whale behaved normally (based on pre-capture behavior and knowledge of normal minke whale behavior), appeared healthy (based on body condition and appearance), and the weather forecast for the next 12 hours was acceptable, the decision was made to start the corralling process.

Before corralling was initiated, the inner aquaculture pen net was lowered to the bottom and the door in the outer net of the pen was opened and tied to the floating ring. The 100 m long E-net was then released from its attachments on the A-net and pulled between two small boats (boat #2 and a second, similarly sized boat) eastwards towards the aquaculture pen (Figure 2). Progress with the E-net was conducted at a slow pace. Once the ends of the E-net reached the nets supporting the aquaculture pen (nets B3 and B2), lines attached to the ends of the E-net were handed to team members on the aquaculture pen so that the ends of the E-net could be manually pulled towards the door opening of the pen’s outer net. During this final stage of corralling, boat engines were turned off to reduce noise. When the water volume between the E-net and the pen door was similar to the volume of the pen itself, the E-net was secured, and the animal was left to find its way through the door into the aquaculture pen. As soon as the whale was observed inside the aquaculture pen, the net section that closes the pen door in the outer net was quickly released from the top and pulled down manually with ropes. Upon closing the door, the inner net was immediately pulled up by the attachment lines. Pulling up the inner net was performed in a controlled manner, utilizing multiple people spread around the aquaculture pen ring. This allowed the net to be pulled up evenly such that no net pockets were created in which the minke whale might get entangled. Once fully contained within the aquaculture pen, the team moved to Phase III.

#### Phase III; Constraining the Whale for Research Procedures

In Phase III of the catch process, the whale was physically controlled in a net hammock for instrumentation, measurement of AEPs (or other physiological measurements), and health monitoring. Observations and recordings of whale behavior and respiration rates were done in 10 min intervals from boat #1, which was docked at the outside of the aquaculture pen ring. After observing the minke whale in the pen for a minimum of 2 hours, and determining that behavior and respiration rates appeared normal, the volume of the aquaculture pen was gradually reduced by first lifting up the inner net to 7 m depth and then pulling a chain of cork floats (roller line) under and across the inner pen net (Figure 3). The roller line was slowly and manually pulled across the pen by six people walking on the floating pen ring. As progress was made, the inner net rolled over the roller line and reduced the water volume around the captured whale (Figure 3). Simultaneously, team members on the raft helped pull the net over the roller. As the roller line approached the opposite side of the ring, a hammock was formed from the remaining net beneath the whale. The whale was finally positioned in the net hammock between the raft and the floating ring of the aquaculture pen; the whale was supported below its body, but still mostly submerged and able to breathe freely. The animal could then be instrumented for health monitoring (ECG, respiration rate and blood draws) and instrumented for AEP measurements (Figure 3). Before release, the whales were instrumented with a finmount satellite tag (Wildlife computers, Splash10-397A single pin design) to monitor post-treatment behavior (i.e. migration and dive behavior).

Post-procedure release of the whale was done by first pulling out the roller line and lowering the inner net to increase the water volume available to the whale. In this manner, the whale could be released back into the full volume of the aquaculture pen within minutes. Observations were recorded for a minimum of 2 hours after release back into the aquaculture pen, after which the inner net was dropped to the sea floor and the door in the outer net opened for the whale to re-enter the catch basin and return to the wild.

### Permits

Permits for this effort were required from the Norwegian Coastal Agency (permit no 2021/6-11/20/40) to anchor nets at sea for six weeks and to divert a shipping route. All nets were clearly marked with floats, buoys and lights at the surface, and navigation warnings were issued. Permits were also required from the Norwegian Fishery Directorate (permit no 22/672 and 23/2507) to catch and release the minke whales. Permits for animal experimentation were given by the Norwegian Animal Research Authority (permit no 19/84343 and 22/241930). Procedures and protocols were also approved by the Institutional Animal Care and Use Committee of the National Marine Mammal Foundation (#15-2019 and #17-2021) with subsequent concurrence by the U.S. Navy Bureau of Medicine and Surgery (NRD 1185). Since the catch-and-release of baleen whales for research purposes is not routine, the permitting process required establishment of a safety protocol for animal handling, an emergency response protocol (if critical situations occurred), and a sedation protocol in case sedation of the whale was deemed necessary by the onsite veterinarian. The sedation protocol was established by a working group of leading marine mammal veterinarians and anesthesiologists.

## Results

### Phase I; Observation and Corralling of the Whale into the Aquaculture Pen

A total of 150 minke whales were sighted and tracked through the CARS over three field seasons - 19 animals in June of 2021, 42 in June of 2022 and 89 in June of 2023. The majority of these animals were adolescent minke whales estimated to be <5 m in length. Most animals approached the CARS from the north, not from the east as expected (Table 1). Sixteen of the 150 animals sighted in phase I (11%) entered the catch basin (Figure 1), of which 10 were contained (phase II), i.e. the whales entered the catch basin and the CARS door was fully closed. The other 6 animals entered the catch basin during the installation phase before the barrier nets were completely in place or escaped back out through the door before it was fully closed. With the 2021/2022 net configuration (Figure 2), many whales escaped the CARS eastwards around the D-nets (Figure 4). When the net configuration was changed in 2023 to prevent this, catch rates increased from 1-2 animals/year in 2021/2022 to 7 animals caught in the catch basin in 2023.

The observed swimming behavior (N=150) in phase I was predominantly calm and normal (CNS 97% of sightings) with the remaining observations being reported as 2% fast vigorous swimming (FVS) and 1% spy hopping (SPY). The average respiration rate in Phase I was 0.6±0.2 breath·min^-1^ (N=61, includes only animals tracked continuously for >5 min). The majority of the sighted minke whales were observed either close to or following the lead nets. Whales were often seen inspecting and probing the nets, but rarely observed physically touching the nets. Our observations clearly demonstrate the ability of minke whales to detect and maneuver around moored nets (black nylon netting), and that it was possible to safely guide this species along such passive net barriers.

### Phase II; Observation and Corralling of the Whale into the Aquaculture Pen

Of the 10 animals contained in the catch basin, 7 animals escaped the basin in phase II, either through gaps between barrier nets, between the barrier nets and the islets or between the barrier nets and the aquaculture pen. Three animals were successfully corralled through the catch basin and contained in the aquaculture pen (Phase III) (Figure 5).

**Figure 5.**
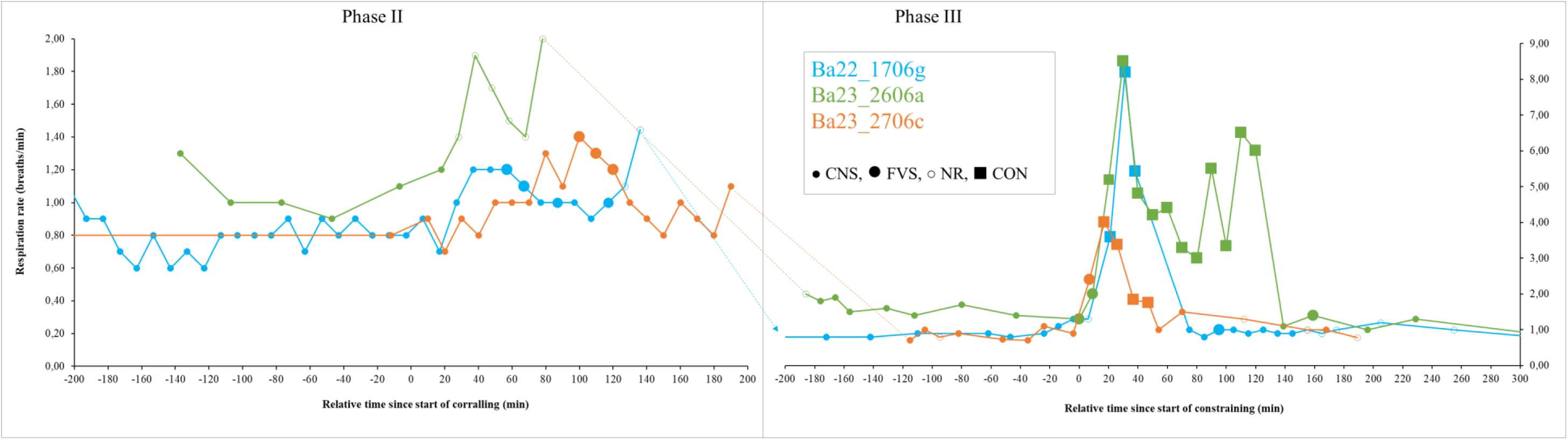
Respiration rate and swim behavior as a function of time in phase II (left panel) and phase III (right panel). The timeline is a relative time (min) since the start of corralling (T=0 min) for phase II and start of reduction of the volume of the aquaculture pen (T=0 min) for phase III. CNS is calm normal swimming, FVS is fast vigorous swimming, NR means behavioral data were not recorded, and CON means the animal was constrained in the net hammock. **Ba22_1706g** was caught at 16:35 (T=-224 min of phase II) on June 17^th^ 2022. Corralling began at 20:19 (T=0 min in phase II) and the whale entered the aquaculture pen at 23:19 (T=-1033 min of Phase III). It was monitored in the pen for 26.8 hours (due to bad weather passing). The process of getting the whale in the hammock was initiated at 16:32 the next day (T=0 min of phase III). The whale was constrained in the hammock at 16:41 (T=9 min of phase III). Due to signs of distress (arching and emesis), the whale was released back into the pen at 17:10 (T=38 min of phase III) before satellite tagging and AEP measurements could be completed. It was observed in the pen for 6.5 hours before it was released back into the ocean at 23:45 (T=433 min of phase III). **Ba23_2606a** was caught at 00:31 (T=-138 min of phase II) on June 26^th^ 2023. Corralling started 02:49 (T=0 min of phase II) and the whale entered the aquaculture pen at 04:11 (T=-200 min of phase III). The constraining process started at 07:21 (T=0 min of phase III), and it was fully constrained in the hammock at 07:41(T=20 min of phase III), released back into the pen at 09:24 (T=123 min of phase III) and back into the ocean at 14:04 (T=403min of phase III). **Ba23_2706c** was caught at 17:18 (T=-1147min of phase II) on June 27^th^ 2023. Corralling started 20:04 (T=-981min of phase II), but corralling failed in bad weather and was therefore paused at 22:30 (T=-835min of phase II). A second attempt was initiated the next day at 12:15 (T=0 min of phase II) and the whale entered the aquaculture pen at 15:28 (T=-132 min of phase III). The constraining process started at 17:40 (T=0min of phase III), it was fully constrained in the hammock at 17:53 (T=13 min of phase III), released back into the pen at 18:30 (T=50 min of phase III) and back into the ocean at 22:30 (T=290 min of phase III).

When the animals entered the catch basin and the door was closed behind them, the animals showed calm normal swimming within the catch basin and average respiration rate of 0.9±0.2 breath·min^-1^ (N=10) in the period before corralling. During corralling of the whales, the swim behavior became slightly faster and more vigorous, and was associated with a small increase in respiration rate, particularly near the end of the corralling where the volume between the opening of the aquaculture pen and the corralling net gets smaller and smaller (Figure 5). Even though the door opening of the outer aquaculture pen net was large (10x15 m), the minke whales hesitated to enter the enclosure of the aquaculture pen, likely due to the large aquaculture ring at the surface of the water under which they had to swim. Nevertheless, the effort demonstrated that minke whales can be actively corralled in a specific direction in a slow and controlled manner by towing a net between two boats (Figure 2).

### Phase III - Constraining the Whale for Research Procedures

The three animals (Ba22_1706g - 3.8 m ?, Ba23_2606a - 4.4 m ♀ and Ba23_2706c - 4.9 m ♀) corralled into the aquaculture pen were all successfully placed in the net hammock for testing and then safely released (Figure 5). Ba22_1706g was held for only 29 minutes before it was released from the hammock due to signs of distress (tachypnea, arching and emesis), in accordance with the project’s animal welfare protocol. As a result, there was insufficient time to complete the AEP measurements and satellite attachment on this animal. Lessons learned from this encounter led to modifications in the handling procedure to provide greater support and control at the axillary region and less constraint of the flukes that improved respiratory efficiency. In Ba23_2606a and Ba23_2706c, AEP measurements of hearing were successfully recorded (Houser et al., 2024) before they were released with a satellite tag attached (Figure 6).

**Figure 6.**
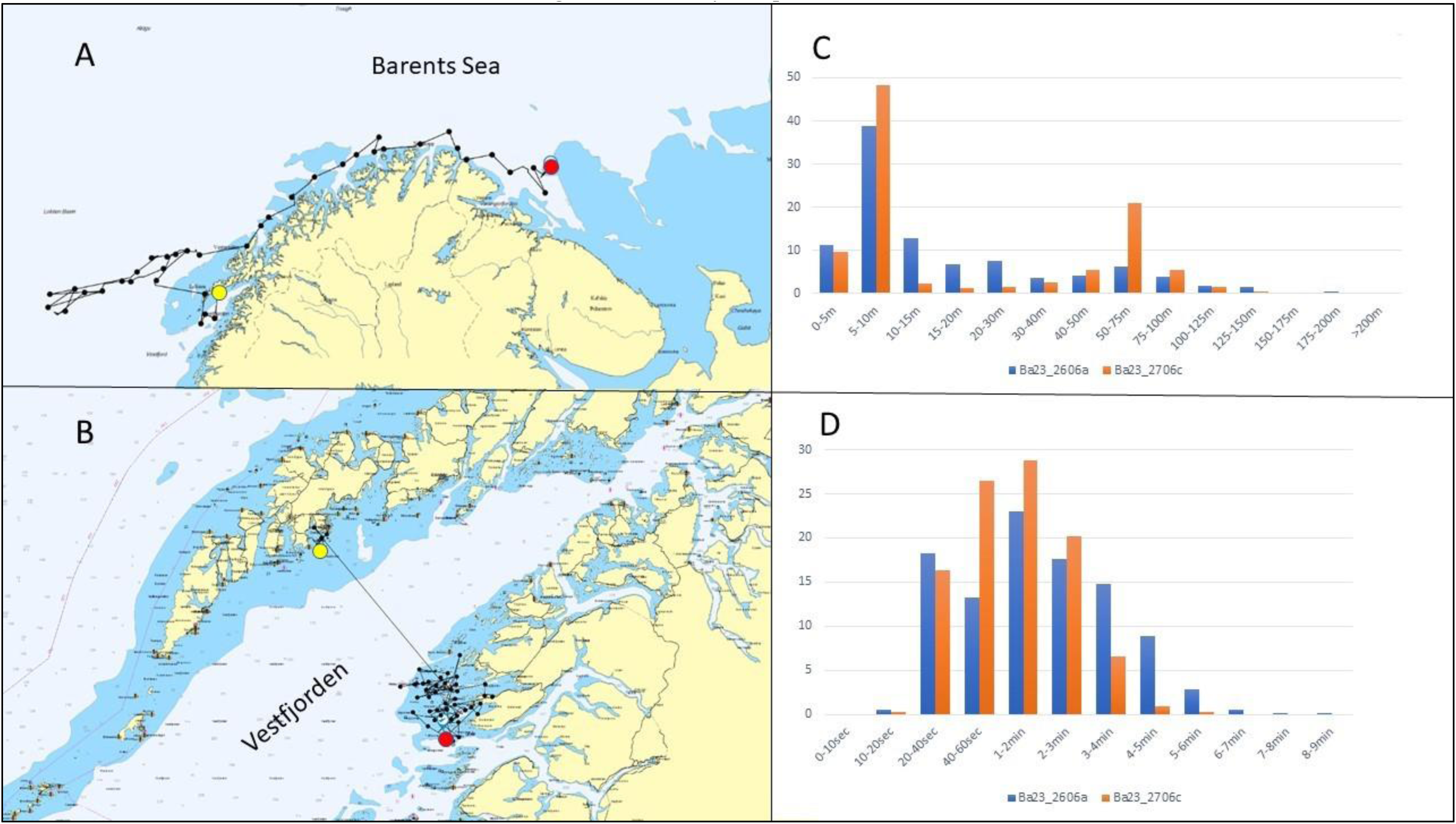
The first two weeks of satellite tracks and dive behavior after release. Panel A and B are the tracks of Ba23_2606a and Ba23_2706c, respectively, with yellow dot at the release position and red dot at the position after 2 weeks. Panel C shows percent dives in depth intervals and Panel D dive duration intervals for both animals for the same period.

As soon as the minke whales entered the fish pen in phase III, they displayed a stereotyped always counterclockwise yet calm swim behavior with a normal respiration rate (Figure 5). When the water volume of the pen was reduced (Figure 3), the animals typically displayed a more erratic vigorous swimming behavior with increasing respiration rates (Figure 5). When they were constrained and touched the net in the hammock, all three animals responded with tonic immobility and tachypnea (Figure 5). Respiration became shallow and more frequent for a few minutes, before it started to decrease again (Figure 5). Upon release back into the aquacultural pen, all three animals immediately returned to the stereotyped counterclockwise, calm swimming behavior with normal respiration rates recorded before handling (Figure 5). After release back into the wild the animals resumed migration and active dive behavior (as recorded by satellite tags), indicating that they did not suffer any long-term effects of the treatment (Figure 6).

## Discussion

A method for the live catch-and-release of minke whales for research purposes has proven feasible, allowing for continuous monitoring during handling, a quick emergency release, and a safe return to the wild. Three whales have been caught and safely released, and initial measurements of their hearing have been made (Houser et al., 2024). Based upon lessons learned over the three years, the method has been steadily modified resulting in progressive improvement in the catch procedure. For example, modifications have been made to the lead net configuration to increase catch rates (Figure 4) and modification of our procedures have been made to improve animal welfare.

Few attempts have been made to capture live mysticete whales for scientific purposes. Prior attempts were either unsuccessful or would be considered unacceptable under modern ethical standards (Vinje, 2022; see introduction for review). Thus, the current project had few relevant studies to rely on while developing our catch methodology. In the Danish pound net fishery, two minke whales have been caught incidentally alive within the past 20 years. These trap nets are tended daily, however the 15-year catch interval makes it unfeasible to be used as a method for planned scientific studies (J. Teilmann, personal observation). Furthermore, the most recent attempt to live-catch baleen whales in Iceland in 2007 (IOGP JIP, 2022) failed because the purse seine net used could not be set quickly enough around the whales (J. Teilmann, personal observation). Learning from these prior approaches, our team determined to use moored stationary lead nets to guide minke whales into a large basin. Once contained in the basin, the whale could be slowly corralled into a smaller enclosure, similar to the Danish pound nets, where it could be lifted to the surface and held for testing.

However, there have also been challenges with the catch-and-release method described here. For example, 7 of the 10 animals caught in the catch basin managed to find gaps between net junctions and escaped. Even though the nets are inspected regularly, the barrier nets are constantly pulled and pushed by ocean currents, tidal forces, and sea waves and thus dynamic gaps might occur. Minke whales orient themselves very precisely around nets (Kot et al., 2012), thereby enabling them to find those gaps and escape if they are given sufficient time. Gaps at net junctions may be mitigated in the future by net placement correction following ROV inspection and by greater net overlap.

### Animal Welfare

The lead and barrier nets used in this study extend to the sea floor and are heavily weighted to eliminate the risk of animal entanglement. Free-hanging ropes and ropes crossing the capture basin were eliminated since fisheries interactions reported for mysticete whales are commonly associated with entanglement in free-hanging ropes, e.g., crab pot lines, and not the net themselves (Song et al., 2010). Other species like humpback whales (*Megaptera novaeangliae)*, killer whales (*Orcinus orca)*, harbour porpoises (*Phocoena phocoena)* and grey seals (*Halichoerus grypus*) were also sighted around the CARS, but as long as the nets were in the intended position, we have observed no incidence involving marine mammal entanglement over the three field seasons conducted thus far. However, in June 2023, prior to being fully operational, unusually strong (full moon) tidal current and strong wind pulled the B1 net 80 m out of position such that one end ended up in deeper water were the lead-line no longer reached the bottom. A minke whale became entangled in the free end of the net and died. This was discovered the next day following CARS repair and ROV inspection of possible damage to nets due to the storm. This animal was not under our care, nor subject to our experimental protocol, nonetheless the catch effort was immediately paused until all procedures were reviewed. After this incident, the anchor points of the nets were re-enforced and the monitoring of the CARS increased in bad weather periods to prevent future incidents due to breakage of the CARS system.

Respiration rates are often used as a diagnostic indication of distress in wild animals (Breed et al., 2019). The average respiration rates of 20 free-ranging minke whales ranged from 0.5 to 1.2 breaths·min^-1^ with a population average of 0.8 breaths·min^-1^ (Øien et al., 2009). More detailed studies looking at respiration rate during different behaviors revealed that during calm normal swimming the respiration rate ranges from 0.5-1.0 breaths·min^-1^, increasing to 1.3-1.5 breaths·min^-1^ during more vigorous swimming (Blix & Folkow, 1995; Folkow & Blix, 1993; Kvadsheim et al., 2017). These observations are consistent with the respiration rates of the whales observed in this study. In confined spaces, surface and respiration rate can increase somewhat without being associated with increased metabolic demand or stress, thus we consider respiration rates between 0.5-1.5 breaths min^-1^ within a 10 min period to be normal in the catch context presented here. Respiration rates between 1.5-2.0 breaths min^-1^ could indicate stress. Our whales exceeded normal respiration and calm swimming behavior only shortly in the last phase of corralling (Figure 5), and what might be considered hyperventilation or tachypnea (>5 breaths/min) was only observed briefly when the whales were fully constrained in the hammock (Figure 5). Thus, based on our observations of respiration rate and swim behavior the last phase of the corralling and being constrained in the hammock seem to be associated with increased stress, but any physiological stress due to handling likely subsided quickly after the stressor was removed. Our animals quickly returned to typical swim behavior and regular respiration rates following release into the aquaculture farm (Figure 5). Furthermore, the whale tracks and the dive data from the satellite tags showed normal behavior following release back into the wild, with no indication of any lasting negative effect from the experiment (Figure 6).

Based on the three catches so far, it seems like this species consistently responds to physical constraint by tonic immobility and tachypnea. Occasionally, arching and emesis were also observed. This implies a need for careful health monitoring of the whales while being held for testing, and release of animals that show signs of distress (decompensation). To minimize distress, our procedures were modified along the course of the work. The last phase of the corralling and the constraining process in the hammock was slowed down and noise minimized (e.g., no machinery nor boat engines are used) to allow the whale to habituate to the smaller volume of water. Intermittently, we also allowed the whale more space in the hammock so that it could freely flex without contacting the net. Finally, we decreased the handling time required for the completion of research procedures (hearing tests and satellite tag attachment) by running these procedures concurrently.

### Future Prospects

Maintaining mysticete whales in aquariums or laboratory facilities is unlikely due to their size, behavior, and feeding requirements. Thus, marine mammalogists would benefit from a field laboratory that is temporarily created in the ocean. Access to temporarily constrained mysticete whales would not only allow for studies of hearing (AEP measurements), but potentially other aspects of sensory physiology, such as sight (Creutzfeldt & Kuhnt, 1973) or tactile senses (Markand, 2020). The potential importance of tactile senses was recently demonstrated in an anatomical study of the Antarctic minke whale where it was proposed that the distributed rigid sensory hairs on the “chin” are used to detect prey and the interface of air and ice (Reichmuth et al., 2022). Other aspects of physiology could also be pursued, such as respirometry (Wahrenbrock et al., 1974), cardiography (Smith & Wahrenbrock, 1974), ultrasonography (Curran & Asher, 1974), tissue (histology) and blood related physiology (hematology, endocrinology, biochemistry panels, lipid analysis, stable isotopes, metabolomics, molecular diagnostics of various diseases), morphometrics (Reidarson et al., 2001), and aspects of animal bioacoustics (Winn et al., 1979). Tagging devices for use on cetaceans have advanced significantly since early tagging attempts by Evans (1974) and Watkins & Schevill (1977). Animal-borne tags are now available that can measure detailed aspects of the behavior and physiology of marine mammals (Andrews et al., 2019; Holton et al., 2021), including heart rate (Goldbogen et al., 2019), blood flow distribution (McKnight et al., 2019), cerebral processes during diving (McKnight et al., 2021), and in the near future, possibly even auditory brainstem response from free ranging animals (Smith et al., 2021). However, such modern tag technology may not be easily deployed remotely, thus, tag attachment will often require access to physically controlled animals, at least for a short period of time.

We acknowledge the complexity of this project, both in terms of field logistics and animal behavior. With our experience in all phases of the project – from handling nets to eliminate gaps at net junctions, corralling and constraining minke whales and measurements of AEP, we will hopefully in the near future provide the first empirical audiogram from a mysticete whale. This will enable us to better understand which sources of anthropogenic noise might impact them and how. In addition, data can potentially be used to guide mysticete auditory weighting functions (e.g., Southall et al., 2019) and validate anatomic hearing models of mysticete whales (e.g., Cranford & Krysl, 2015).

## Acknowledgments

This project was funded by the US Subcommittee on Ocean Science and Technology (SOST) Interagency Working Group on Ocean Sound and Marine Life, which is a partnership between the Office of Naval Research (contract # N0001420C2022), Chief of Naval Operations N45 (the Navy Living Marine Resources program, contract # N3943019C2167), the Bureau of Ocean Energy Management (contract # N0001420C2022), the National Oceanic and Atmospheric Administration (contract #UCAR SUBAWD002120), and the Marine Mammal Commission (contract #MMC 19167). We are very thankful to all team members who participated in the field effort (H Axelsen, J Balle, K Bøkenes, D Gaudet, C Fisher, E Franks, H Førde, G Goya, SR Hansen, E Hofoss, TV Johnsen, K Kleivane, A Kumar, JA Marcussen, R Otnes, R Roland, D Schreher, T Sivertsen, M Shoemaker, T Tjomsland and M Weise). Special thanks to Kurt Svendsen and Henrik Svendsen and their crew at IsQueen AS for all logistical support, Selstad and Mørenot AS for their services, the late Tommy Sivertsen and his son, Thomas Sivertsen, for sharing their local knowledge, and the crews of MS Andopsvaeringen, MS Ballstadvaeringen, MS Einar Erlend, MS Roger and MS Kurt Sr. for assistance in handling more than 40 tons of nets. The sedation protocol was developed by a working group consisting of Drs. J Bailey, C Harms, D Houser, M Moore and RA Ølberg.

All authors declare no competing interest.

## Notes

https://www.ffi.no/en/research/the-minke-whale-hearing-project

https://www.nmmf.org/our-work/biologic-bioacoustic-research/minke-whale-hearing/

### Competing Interest Statement

The authors have declared no competing interest.

